# Guiding G protein signaling by target enhancement of GPCRs

**DOI:** 10.1101/2025.02.06.636923

**Authors:** Brix Mücher, Aida Garrido-Charles, Lukas Cyganek, Tobias Bruegmann, Deniz Dalkara, Ida Siveke, Stefan Herlitze

## Abstract

Activation of G protein coupled receptors coupling to the G_i/o_ pathway leads to the activation of G protein-activated inward rectifier potassium channels (GIRK) in a fast membrane-delimited manner in excitable cells. Activation of GIRK causes the hyperpolarization of the cell membrane, where hyperpolarization is dependent on te availability of G_i/o_ coupled GPCRs and GIRK. In particular, in optogenetic and chemogenetic experiments neuronal silencing depends on downstream targets of G_i/o_-coupled GPCRs. To selectively enhance G_i/o_ mediated GIRK currents, we created expression cassettes consisting of a homomer forming GIRK subunit and various light-activated G_i/o_-coupled GPCRs (Melanopsin, Long-wave-sensitive opsin 1, Parapinopsin or Opsin 7b). We demonstrate that light-activation of the GIRK/GPCR constructs induces robust GIRK currents in human embryonic kidney 293 cells, cardiomyocytes and cerebellar Purkinje cells and changes the net effect of G protein signaling of the promiscuous Opn4L from a G_q/11_ mediated excitation towards an G_i/o_ mediated inhibition. Thus, our tools enhance target selectivity and improve optogenetic control of the G_i/o_ pathway by light in excitable cells.

## Introduction

Guanine nucleotide binding protein (G-protein)-coupled receptors (GPCRs) are an ancient and highly diversified family of receptors that convert extracellular chemical or physical stimuli such as light into diverse multi-layered intracellular signaling cascades. The protein superfamily of GPCRs is found in protozoa^1,2^, fungi^3,4^, plants^5–7^ and throughout all phyla of the animal kingdom with around 800 genes discovered in humans^8,9^. GPCRs consist of seven transmembrane domains (TMD), with an extracellular N- and an intracellular C-terminus^10–13^. Ligand specificity is majorly determined by extracellular regions including binding pockets within the TMD^13–15^, while a GPCR interacts with different classes of G proteins at its intracellular domain^13,16–20^. G proteins consist of a guanosine triphosphate (GTP) binding Gα subunit and a heterodimeric Gβγ complex. Four main families of Gα subunits can be distinguished: the adenylate cyclase-stimulating Gα protein (Gα_s_) family (G_sS&XL_-^21–28^ and G_olf_-subunits^29,30^), the adenylate cyclase-inhibiting Gα protein (Gα_i/o_) family (G_i1-3_-^23,27,28,31–34^, G_o1&2_-^35–39^, G_t1&2_-^40–42^, G_z_-^43,44^ and G_gust_^45,46^-subunits), the Gα_q_ family (G_q_-^47^, G_11_^48,49^-, G_14_-^49,50^ and G_15/16_-subunits^51,52^), which activates phospholipase C (PLC), and the Gα_12_ family (G_12&13_^49,53–55^-subunits), which activates rho GTPases. As mentioned above Gβ forms dimeric complexes with Gγ, which once released from the heterotrimeric G-protein complex, mediate a wide range of effects, such as the recycling of GPCRs, the activation or inhibition of various ion channels (i.e. G protein-activated inward rectifier potassium channel (GIRK)^56–61^, voltage-gated calcium channel (VGCC)^62–64^ and transient receptor potential channel^65,66^) or the interaction with canonical Gα targets like PLC^67–69^ or adenylyl cyclase^70–72^.

Considering the plethora of downstream targets for GPCRs, the diversity of ligands and cellular functions and their presence in varying composition in almost all cell types and tissues of animals, it is not surprising that GPCRs can also play a major role during the pathogenesis and treatment of psychiatric disorders, such as schizophrenia^73^ or depressive disorder^74^. Furthermore, GPCRs are implicated during neurodegenerative diseases, including Alzheimer’s^75,76^ or Huntington^77,78^, as well as metabolic disorders^79^ or cancer^80–82^. Consequently, the prevalence of GPCRs in disease highlights the importance of unraveling their multidimensional signaling cascades and cellular targets.

GPCRs signal in a cellular and temporal specific manner dependent on intracellular signaling targets. One important target of G_i/o_-coupled GPCRs in excitable cells are GIRKs. These channels are activated by Gβγ subunits in a membrane-delimited manner causing an efflux of potassium and subsequent membrane hyperpolarization. Four different GIRK subunits have been identified (GIRK1 (Kir3.1), GIRK2 (Kir3.2), GIRK3 (Kir3.3), and GIRK4 (Kir3.4)). GIRKs consist of four subunits and are often found as heterotetramers in heart (most common: GIRK1, 4) and brain (most common: GIRK1, 2). Functional GIRK2 and 4 homomeric channels^83–85^ have been described, although they display smaller currents than GIRK1-containing channels^86^. GIRK1 lacks crucial export motifs for the endoplasmic reticulum and GIRK3 redirects channels to lysosomes^87^. Functional homomeric channels for GIRK1 subunits have not been detected. The point mutation F137S^88,89^ in GIRK1, however, can form functional homomers, likely by eliminating the large non-conserved phenylalanine from its pore.

Light-activated GPCRs allow for an optogenetic approach to study and modulate cell-type specific effects of GPCR signaling pathways with high temporal and spatial precision^90–92^. GIRK, as a direct target in the GPCR signal transduction, is used to characterize G protein specificity and the affinity and action spectrum of ligand- and light-activated GPCRs. More importantly, silencing of excitable cells using chemogenetic and optogenetic strategies (DREADDs or OptoXRs) relies on the endogenous expression of somatodendritically localized GIRK channels and presynaptically localized VGCCs^93^ in neurons and in cardiomyocytes within the sinus node and atria. Somatodendritic localization allows GIRK to regulate postsynaptic excitability and the firing rate of neurons in response to transmitter release. Here, hyperpolarization is limited by the level of GIRK channel expression and its subunit composition.

To guarantee sufficient G_i/o_ targets in excitable cell types, which, for example express low GIRK currents, we established an adeno-associated virus (AAV) based, single expression cassette approach, where we co-express a homomer forming GIRK channel subunit together with different G_i/o_-coupled light-activated GPCRs. First, we tested different GIRK channel isoforms and mutants for their potential to form homomeric channels in human embryonic kidney 293 (HEK293) cells. Second, the GIRK1 subunit pore mutant F137S^88,89^ was selected and characterized in combination with four different light-activated GPCRs (the multistable G_q_ and G_i_ coupling mouse melanopsin^94–97^ (Opn4L), the G_t-_coupled human long wavelength opsin^98–101^ (Opn1LW), the bistable G_i_-coupled lamprey parapinopsin^102–107^ (ppOpn) and the constitutively active G_i_-coupled zebrafish novopsin-4 (Opn7b)^108,109^). Third, we validated the functionality and potency of the GIRK-Opn4L construct to study and drive G protein/GIRK-based inhibition in human induced pluripotent stem cell derived cardiomyocytes (hiPSC-CMs), a premature model of ventricular cardiomyocytes with spontaneous beating and in cerebellar Purkinje cells (PC). Furthermore, we demonstrate a shift in excitability of PC firing by inducing G_i/o_ mediated GIRK channel activation using the co-expression of the promiscuous Opn4L and GIRK1^F137S^. In summary, our tools, which comprise a light-activated GPCR and an ion channel target not only highlight the importance of intracellular G protein targets, but also guide GPCR signaling and provide improved optogenetic control over the G_i/o_ pathway by light.

## Results

### Gβγ activated GIRK homomultimers for co-expression with G_i/o_-coupled GPCRs

Since co-expression of optogenetic tools with sensors or target proteins using AAVs is limited by the DNA insertion capacity of the AAV, we needed to identify GIRK channels that display robust light induced currents when composed of a single subunit. The GIRK subunit mutants GIRK1^F137S^ and GIRK4^S143T^, have been described to form functional, ion conducting GIRK homomers^88,89^, such as the naturally occurring homomultimer forming subunit GIRK2. GIRK subunits reveal a high degree of homology within their pore-forming intramembrane region. However, GIRK1 has a phenylalanine (F) at position 137, while the other subunits have a conserved serine (S) residue at the corresponding position (Fig. 1a red; other differences within the pore P131, A142 for GIRK1 and F154 for GIRK4 are highlighted in grey). We compared these subunits to GIRK1 monomers and GIRK1, 2 heteromers in HEK293 cells stably expressing mouse Opn4L (activated at 470 nm). Opn4L couples to the G_i/o_ pathway^95^ and induces GIRK1, 2 currents^96^. Currents differed depending on the expressed GIRK subunits (Fig. 1c). Expression of the pore mutants as well as the heteromeric expression of GIRK resulted in significantly larger currents than controls (Fig. 1c). Expression of GIRK1^F137S^ allowed for large light-induced potassium currents (Fig. 1b, c), which were not seen with the wild-type GIRK1 subunit (Fig. 1b, c). The GIRK4 channel mutant GIRK4^S143T^ also gave rise to large light-induced potassium currents (Fig. 1b, c). Overexpression of the GIRK1^F137S^ or GIRK4^S143T^ subunit allowed for larger light-induced potassium currents than GIRK2 homomeric channels (Fig. 1b, c). The heteromeric expression of GIRK1 and GIRK2 significantly increased the inducible GIRK current compared to GIRK2 expression (Fig. 1b, c). Channel activation times differed between the different GIRK channels. Light-induced GIRK1^F137S^ homomultimer currents were activated slightly faster than GIRK4^S143T^ currents (Fig. 1d), but channel deactivation times were comparable between all tested channels (Fig. 1e).

**Figure 1:**
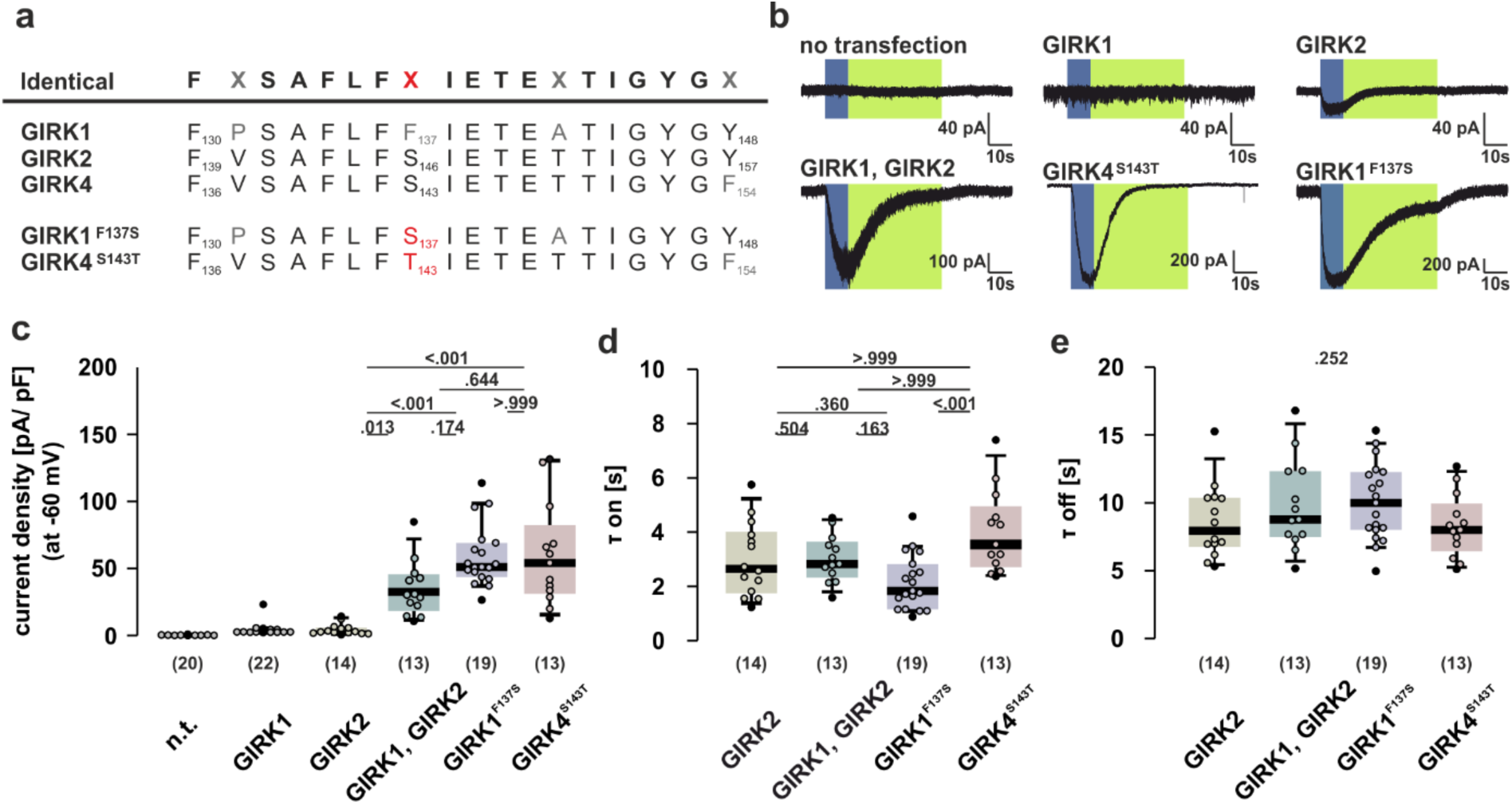
Comparison of light-activated currents mediated by GIRK homo-/ heteromers in HEK293 cells stably expressing Opn4L. **a)** Alignment of the amino acids of the pore region of GIRK subunits with introduced mutations (red) and differences (light grey) from the consensus sequence (black). **b)** Exemplary traces of light-induced currents of GIRK expressing HEK293 cells constitutively expressing Opn4L. **c)** Current density, **d)** τ on, and **e)** τ off from GIRK currents elicited during 10 s blue light stimulation of Opn4L-HEK293 cells expressing GIRK1 (grey), GIRK2 (beige), GIRK1, 2 (green), GIRK1^F137S^ (blue), or GIRK4^S143T^ (red). Boxplots depict median, 10^th^, 25^th^, 75^th^, and 90^th^ percentiles. Outliers are shown as black circles while other cells are displayed as colored circles.

### Establishing the light-modulated GIRK - G_i/o_-coupled GPCR co-expression cassettes in HEK293 cells

To test feasibility and characterize functionality of the GIRK-OpnX co-expression cassette, we engineered a GIRK1^F137S^ p2A Opn4L.eGFP (GIRK-Opn4L) construct and characterized this construct in HEK293 cells. Membrane expression of this construct suggests functional co-expression of GIRK and Opn4L.eGFP using the p2A peptide (Fig. 2a_1_). Blue light stimulation (470 nm, 10 s) of GIRK-Opn4L results in an activation of GIRK1^F137S^ and a subsequently increasing potassium current (Fig. 2b_1_, τ_on_ ≈ 1.5 s; Fig. 2c), while green/yellow light (560 nm) leads to a deactivation and reduction of current (Fig. 2b_1_, τ_off_ ≈ 9.9 s Fig. 2d).

**Figure 2:**
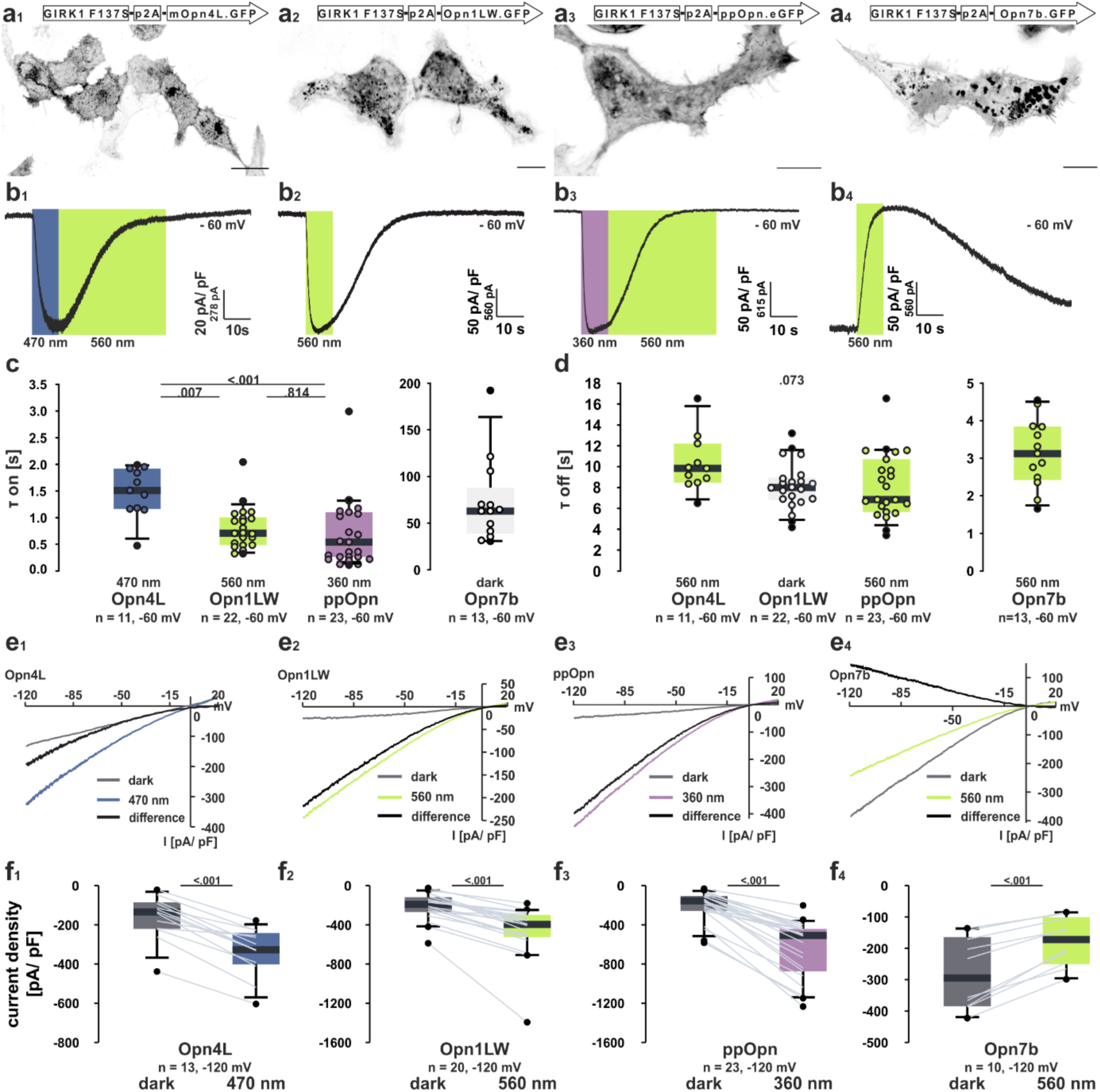
Characterization of the GIRK-OpnX co-expression cassette in HEK293 cells. GIRK1^F137S^ p2A **a_1_)** Opn4L **a_2_)** Opn1LW, **a_3_)** ppOpn and **a_4_)** Opn7b fused to eGFP expressed in HEK293 cells. Scale bars represent 20 µm. GIRK-OpnX currents are induced by activation of **b_1_)** Opn4L (blue, 470 nm), **b_2_)** Opn1LW (lime, 560 nm), and **b_3_)** ppOpn (UV, 360 nm), while stimulation of **b_4_)** Opn7b (lime, 560 nm) reduces GIRK currents. Currents can be deactivated by light stimulation of **b_1_)** Opn4L and **b_3_)** ppOpn with a yellowish-green light (560 nm), while **b_2_)** Opn1LW and **b_4_)** Opn7b induced currents return to basal levels in the dark. **c)** Activation and **d)** deactivation times of GIRK currents in cells expressing GIRK-OpnX. **e)** Exemplary IV-curves illustrating GIRK currents elicited at different membrane voltages in the dark (grey) or during light stimulation of **e_1_)** Opn4L (470 nm), **e_2_)** Opn1LW (560 nm), **e_3_)** ppOpn (360 nm), and **e_4_)** Opn7b (560 nm). The difference between light-activated and basal currents in the dark (black) is also displayed. Current density of **f_1_)** GIRK-Opn4L, **f_2_)** GIRK-Opn1LW, **f_3_)** GIRK-ppOpn, and **f_4_)** GIRK-Opn7b expressing cells at −120 mV in the dark (grey) or when stimulated with light of the respective activation wavelength (color). Boxplots depict median, 10^th^, 25^th^, 75^th^, and 90^th^ percentiles. Individual cells are indicated by grey lines, while outliers are shown as black circles.

Since we could successfully control light-induced GIRK currents using the GIRK-Opn4L construct, we engineered further GIRK-OpnX expression cassettes. We chose the G_i/o_-coupled human long-wave-sensitive opsin 1 (Opn1LW, Fig. 2a_2_, 2b_2_), the bistable lamprey parapinopsin (ppOpn, bistable UV pigment from *Lethenteron camtschaticum*, Fig. 2a_3_, 2b_3_), and the reverse photoreceptor Novopsin-4 (Opn7b, *Danio rerio*, Fig. 2a_4_, 2b_4_). Light-activation of GIRK-Opn1LW.eGFP (560 nm, Fig. 2b_2_, τ_on_ ≈ 0.7 s, Fig. 2c) and GIRK-ppOpn.eGFP (360 mV, Fig. 2b_3_, τ_on_ ≈ 0.5 s, Fig. 2c) led to robust, fast increases in GIRK currents, which declined to baseline levels once the light was switched off (Opn1LW, τ_off_ ≈ 8 s Fig. 2d) or switched to green/ yellow light (ppOpn, 560 nm, Fig. 2b, τ_off_ ≈ 6.8 s Fig. 2d). Comparison of activation and deactivation times revealed the strictly G_i/o_-coupled opsins Opn1LW and ppOpn to be faster in activating GIRK1^F137S^ when compared to Opn4L (Fig. 2c), while deactivation times were comparable (Fig. 2d). In contrast to the opsins described above, light-induced deactivation of GIRK-Opn7b.eGFP (560 nm, Fig. 2b_4_, τ_off_ ≈ 2.9, Fig. 2d, τ_on_ ≈ 62.9 s Fig. 2c) decreased the continuous GIRK current, which recovered to baseline levels in the dark. Exemplary current-voltage characteristics (Fig. 2e_1-4_) illustrate the large inward rectifying current that could be elicited by the activation of GIRK-OpnX constructs by light (Fig. 2f_1-4_).

To analyze changes in the constitutive activity of the GIRK-OpnX expression constructs, we compared the baseline and light-induced potassium currents in GIRK-OpnX expressing HEK293 cells between currents measured in solutions with high and low potassium concentrations. Switching from high to low potassium concentration leads to a negative shift in reversal potential close to the holding potential of −60 mV, thereby drastically reducing the inward current. Light induced currents were always larger than resting potassium currents for Opn4L (Fig. 3a_1_), Opn1LW (Fig. 3a_2_) and ppOpn (Fig. 3a_3_). We found that GIRK-Opn4L and GIRK-Opn1LW reveal a relative mean constitutive channel activity of 35% and 31% in high potassium conditions respectively (Fig. 3b_1_). GIRK-ppOpn shows a lower mean basal current of only 20% of baseline activity relative to total current under high potassium concentration (Fig. 3b_1_), resulting in a significantly larger percentage of light inducible current (Fig. 3b_2_). Activity of the reverse photoreceptor GIRK-Opn7b can be reduced by 43% (mean) of the total current by light (Fig. 3b_3_). In summary, each respective co-expression cassette of GIRK-Opn4L, GIRK-Opn1LW, GIRK-ppOpn or GIRK-Opn7b can be used to induce and activate or deactivate GIRK currents by light.

**Figure 3:**
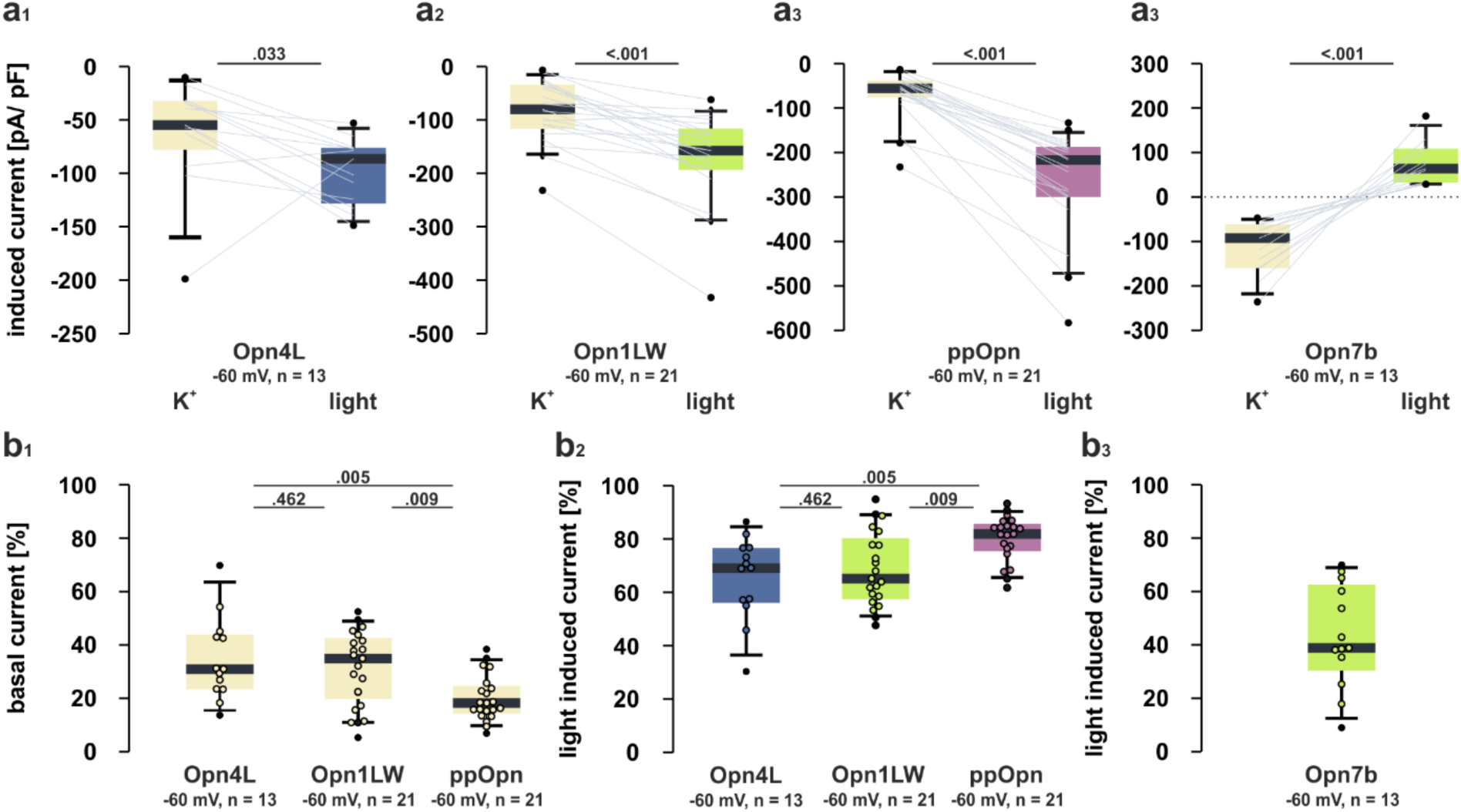
Comparison of constitutive (basal) and light-induced currents in GIRK-OpnX expressing HEK293 cells. **a)** Current density at HEK293 membranes expressing **a_1_)** GIRK1^F137S^-Opn4L, **a_2_)** GIRK1^F137S^-Opn1LW, **a_3_)** GIRK1^F137S^-ppOpn, or **a_4_)** GIRK1^F137S^-Opn7b. Currents induced by a switch in external potassium concentration in the dark (beige), are lower than currents induced by light under high external potassium conditions (blue, green, purple). Percentage of **b_1_)** constitutive or **b_2_)** light-induced current for GIRK1^F137S^-Opn4L, GIRK1^F137S^-Opn1LW and GIRK1^F137S^-ppOpn expressing cells. **b_3_)** Light-induced reduction in current density by GIRK1^F137S^-Opn7b (the current density is at its maximum under high potassium conditions). Percentages in b) from recordings depicted in a). Boxplots show median, 10^th^, 25^th^, 75^th^ and 90^th^ percentiles. Outliers are shown as black circles, while other cells are displayed as colored circles or grey lines.

### Measurement of GIRK-OpnX co-expression cassettes in cerebellar Purkinje cells

Cerebellar Purkinje cells express GIRK channels in moderate levels making it difficult to recruit endogenous GIRK channel currents for G_i/o_ mediated somatodendritic inhibition using chemical ligand- or light-activated G_i/o_ coupled receptors^110,111^. Therefore, to gain an understanding of light-induced GIRK channel currents in a cell-type with moderate levels of endogenous inwardly rectifying potassium current, we expressed Opn4L, Opn1LW, ppOpn or Opn7b and compared light-induced currents to GIRK-Opn4L, GIRK-Opn1LW, GIRK-ppOpn or GIRK-Opn7b induced currents. We performed voltage-clamp recordings in cerebellar Purkinje cells and monitored changes in membrane currents during voltage ramps between −120 to −60 mV before and during light application (Fig. 4a-d). We observed an increase in inward current between −120 to −60 mV for the light-activated GIRK-Opn1LW, GIRK-Opn4L and GIRK-ppOpn expressing but not for Opn1LW, Opn4L or ppOpn expressing PCs (Fig. 4e-g). The constitutively active opsin Opn7b in combination with GIRK but not Opn7b alone reduced the membrane current during light application (Figure 4h).

**Figure 4:**
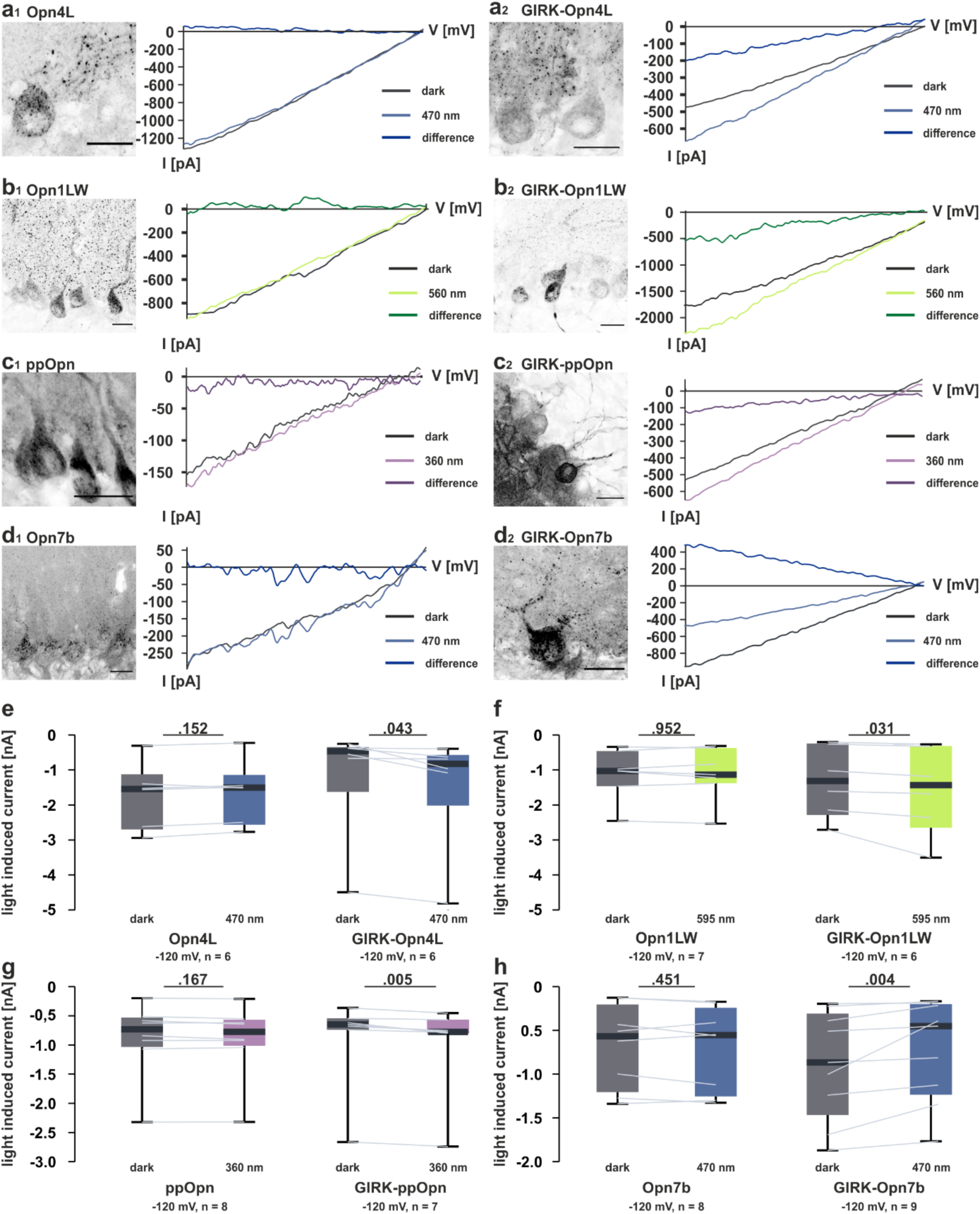
Comparison of light-induced Opn and GIRK1^F137S^-OpnX currents in cerebellar Purkinje cells. **a-d)** Exemplary IV-curves illustrate the current response of cerebellar Purkinje cells to voltage changes between −120 to −60 mV for dark adapted (grey trace) and light-stimulated (colored trace) cells. For **a_1_)** Opn4L, **b_1_)** Opn1LW, **C_1_)** ppOpn and **d_1_)** Opn7b no difference in Purkinje cell current is seen upon light stimulation, while **a_2_)** GIRK1^F137S^-Opn4L, **b_2_)** GIRK1^F137S^-Opn1LW, **c_2_)** GIRK1^F137S^-ppOpn and **d_2_)** GIRK1^F137S^-Opn7b induce a change in Purkinje cell current when stimulated. Scale bars represent 20 µm. **e-h)** Differences in light-induced currents at −120 mV between dark state (grey) and light-activated states (colored) of **e)** Opn4L/ GIRK1^F137S^-Opn4L, **f)** Opn1LW/ GIRK1^F137S^-Opn4L, **g)** ppOpn/ GIRK1^F137S^-Opn4L and, **h)** Opn7b/ GIRK1^F137S^-Opn4L expressing Purkinje cells. Boxplots depict median, 10^th^, 25^th^, 75^th^ and 90^th^ percentiles. Outliers are shown as black circles, while other cells are displayed as colored circles or grey lines.

### Co-expression of GIRK1^F137S^ and Opn4L shifts light-induced excitation towards inhibition

Melanopsin activates various G protein signaling pathways including the G_i/o_ and G_q/11_ pathway^96^. We could previously show that the activation of Opn4L induces depolarization and increases action potential (AP) firing in cerebellar PCs, indicating a dominant G_q/11_ pathway activation^96^. In HEK293 cells expressing GIRK1, 2 subunits together with Opn4L, both G_i/o_ and G_q/11_ pathways can be activated by light at the same time (Fig. 5). Supplementary expression of GIRK1^F137S^ using the co-expression cassette shifted the G_q/11_/G_i/o_ balance from G_q/11_ mediated intracellular Ca^2+^ increase to G_i/o_ mediated GIRK channel activation (Fig. 5).

**Figure 5:**
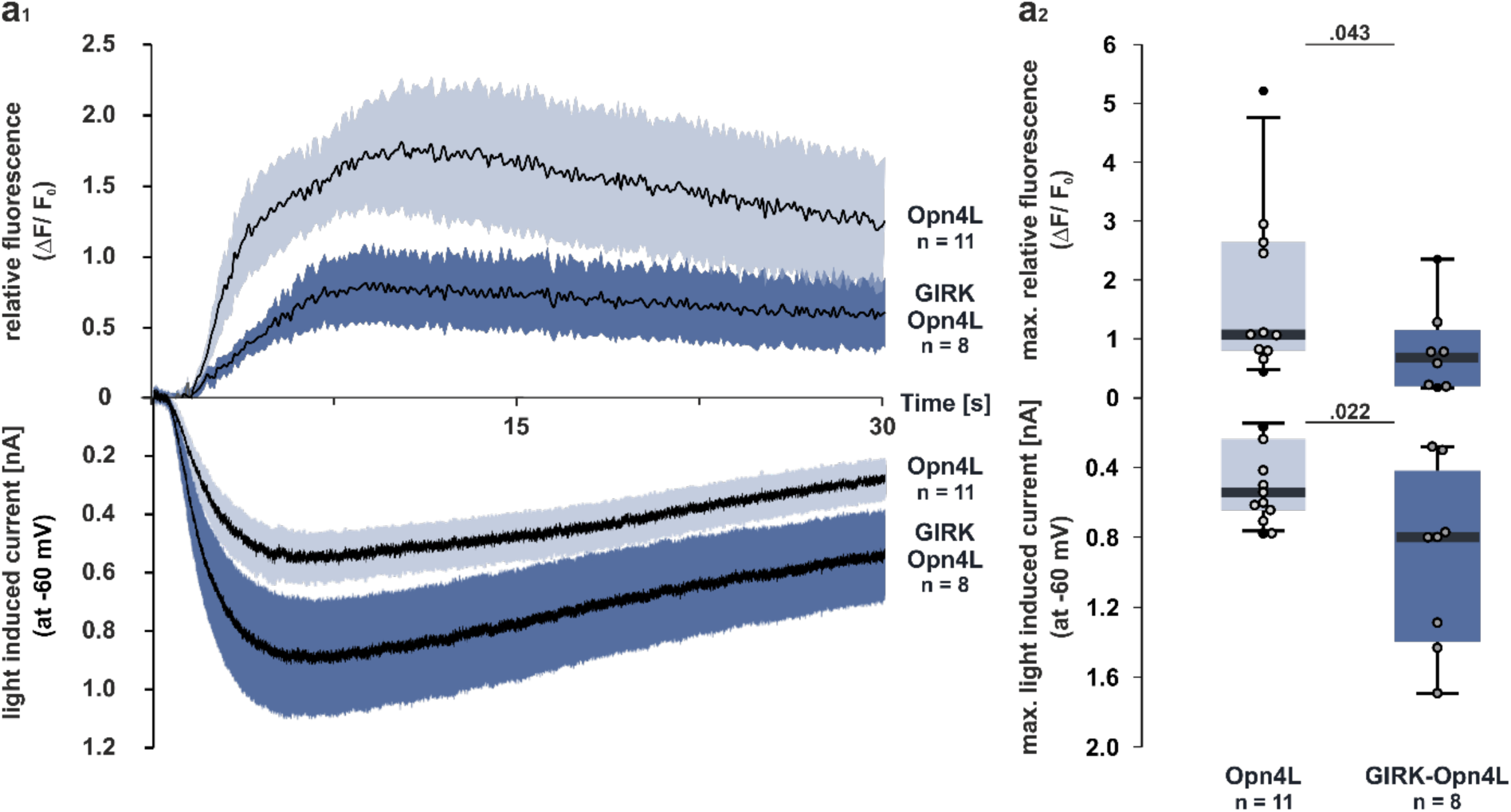
Comparison of Opn4L induced G_q/11_-mediated calcium and G_i/o_-activated GIRK dynamics in HEK293 cells stably expressing GIRK1, 2. Cells were transfected with either GIRK1^F137S^-Opn4L (dark blue) or Opn4L (light blue) and GCaMP6f. **a_1_)** Cells respond to blue light (470 nm, constant stimulation) with an increase in intracellular calcium (upper traces) and potassium (lower traces). Data are presented as mean (black) alongside its standard error (light/ dark blue). **a_2_)** Maximum fluorescence reveals significantly lower intracellular calcium levels for the co-expression of GIRK1^F137S^ (dark blue) compared to Opn4L expression alone (light grey), while the peak light induced current displayed is significantly increased when additional GIRK1^F137S^ subunits are co-expressed (dark blue) compared to the stimulation of Opn4L alone (light blue). Boxplots depict median, 10^th^, 25^th^, 75^th^ and 90^th^ percentiles, outliers are shown as black circles and other cells are displayed as blue circles.

We therefore examined if the GIRK-Opn4L expression cassette shifts the pathway predominance of Opn4L from a G_q/11_ mediated excitation to G_i/o_ and GIRK mediated inhibition in cerebellar PCs. We analyzed whether light-activated GIRK-Opn4L would decrease the AP firing rate and would act opposing to the light-induced excitation mediated by Opn4L alone. As demonstrated in Figure 6 the light-induced activation of Opn4L increases the spontaneous AP firing rate of PCs (Fig. 6b_1_, c), as well as the evoked firing rate (Fig. 6d_1_, e), while light-induced activation of GIRK-Opn4L decreases spontaneous and evoked firing (Fig. 6b_2_, c, f). The modulated excitability due to supplemental expression of GIRK is further revealed by the change in rheobase upon light stimulation (Figure 6h). In addition, a decrease in input resistance, indicating more open ion channels at the cell membrane, is only observed for GIRK-Opn4L but not Opn4L expression (Fig. 6j).

**Figure 6:**
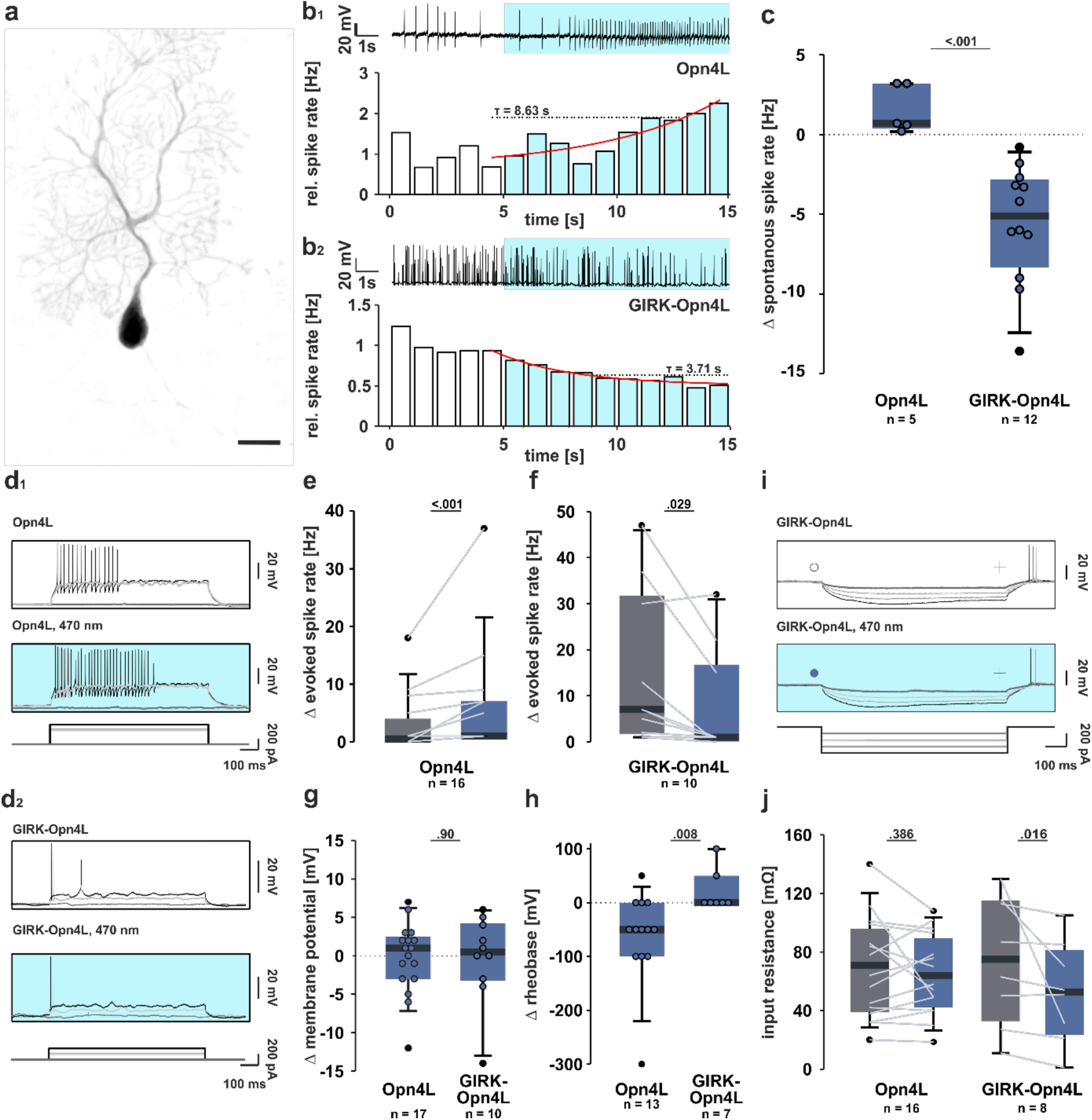
Co-expression of GIRK1^F137S^ shifts Opn4L induced excitation to inhibition in cerebellar Purkinje cells. **a)** A cerebellar Purkinje cell expressing GIRK1^F137S^-Opn4L; the scale bar represents 20 µm. **b_2_)** GIRK1^F137S^-Opn4L expression results in a reduced spontanous spike rate of cerebellar Purkinje cells in response to light (470 nm, blue), while **b_1_)** Opn4L expressing cells respond with an increased action potential firing rate. **c)** Changes in the spontaneous spike rate are significantly different for Opn4L and GIRK^F137S^-Opn4L expression. **d_1_, e)** Evoked spikes of Purkinje cells increase upon light stimulation (470 nm, blue) of Opn4L, but **d_2_, f)** decrease during the light stimulation (470 nm, blue) of GIRK^F137S^-Opn4L expressing cells if compared to spikes recorded in the dark (grey). **g)** The membrane potential of Purkinje cells remained unchanged but **h)** the rheobase was significantly increased for GIRK^F137S^-Opn4L expression compared to Opn4L expression alone (applied in 50 pA steps). **i, j)** No change in input resistance was recorded during light stimulation (470 nm, blue) of Opn4L expressing cells in comparison to cells in the dark (grey), but excitation of GIRK^F137S^-Opn4L (470 nm, blue) expressing cells resulted in a significantly lower input resistance. Boxplots depict median, 10th, 25th, 75th and 90th percentiles, outliers are shown as black circles, while other cells are displayed as colored circles or grey lines for comparative measurements.

### Co-expression of GIRK1^F137S^ and Opn4L decreases spontaneous beating in cardiomyocytes

GIRK channel activation is one key part of the parasympathetic regulation of heart activity. In pacemaking cells of the sinus node and atrioventricular node, as well as in atrial cardiomyocytes, activation of the muscarine receptor type 2 induces increased GIRK currents leading to reduced spontaneous beating rate and shortened action potential duration^112^. Stem cell derived cardiomyocytes enable investigations of the regulation of automaticity by GPCRs since they spontaneously beat due to their fetal maturation state. We have shown that mouse Opn4 leads to a light-induced increase in the spontaneous beating rate due to enhanced calcium clock machinery in transgenic murine embryonic stem cell derived cardiomyocytes^113^. In contrast, in the adult heart, mouse Opn4 suppressed spontaneous beating proving its promiscuity and the more pronounced G_i_ signaling in the adult heart^114^. Together, both observations demonstrate the ability of Opn4 to activate both G_i_ and G_q_ pathways in cardiomyocytes and that in fetal, premature cardiomocytes the G_q_ signaling effect prevails. To demonstrate that, also in cardiomyocytes, overexpression of GIRK alongside Opn4L can switch the light effect towards G_i_ protein mediated decrease in spontaneous beating, we expressed Opn4L alone or together with GIRK1^F137S^ in hiPSC-CMs resulting in membrane restricted GFP signals (Fig. 7A and B). Illumination with blue light (470 nm, 5 s, 2.7 mW/mm^2^) induced a significant reduction in the spontaneous beating rate of cardiomyocytes expressing GIRK-Opn4L but not Opn4L alone (Fig. 7c).

**Figure 7:**
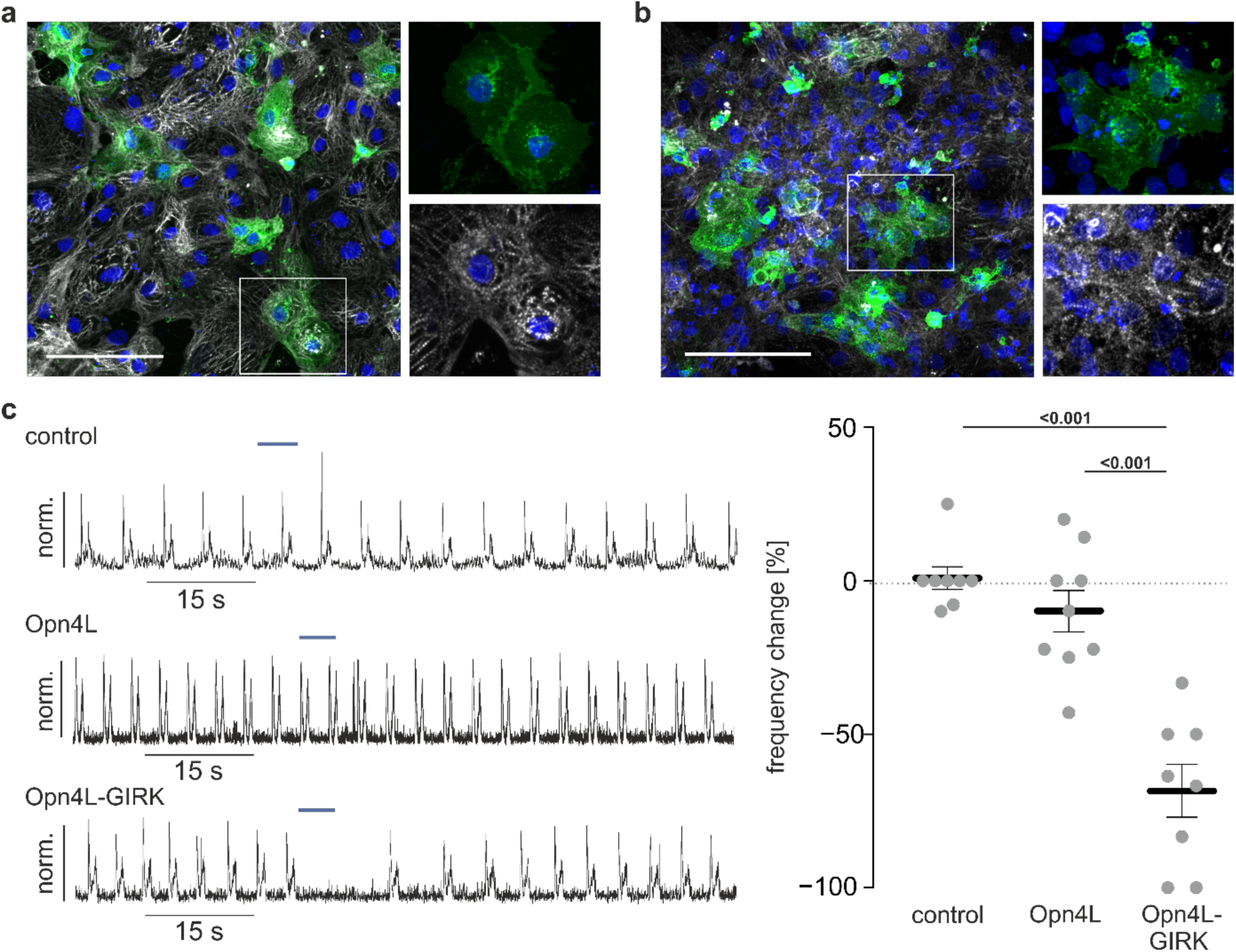
Co-expression of GIRK1^F137S^ enables Opn4L mediated reduction of spontaneous beating rate in cardiomyocytes. a-b) Confocal images of cardiac cell clusters consisting of hiPSC-CMs cardiomyocytes identified by cardiac troponin T (white) and stained for nuclei with DAPI (blue) overexpressing Opn4L.eGFP (green, A) or GIRK-Opn4L.eGFP (green, B). Scale bar is 100 µm. c) Representative traces and aggregated data of spontaneous beating of control cardiac cell clusters (top), cardiac cell clusters expressing GIRK-Opn4L.eGFP (middle) or Opn4L.eGFP (bottom). Circles represent the result from one cardiac cell cluster. The mean is shown with the standard error of the mean as error bars.

## Discussion

### Single vector based optogenetic tools to guide GPCR function

AAVs have emerged as leading vectors for gene therapy and *in vivo* research^115^. They show wide tropisms and catalyze a low adaptive immune response in humans, but the major limitation asides persistence of AAVs is their relatively small packaging capacity, which is considered to be around 4.7 kilobases but depends on the AAV capsid used^115–118^. To reduce the size of our G_i/o_ co-expression cassette we incorporated a shortened expression cassette^119^ and explored the possibility to use GIRK subunits that assemble to form homomeric channels. As demonstrated before pore mutants of both GIRK1 and GIRK4 allowed for the formation of functional GIRK homomers. In comparison to the expression of GIRK1 or GIRK2 the light-induced Opn4L mediated GIRK1^F137S^ and GIRK4^S143T^ currents were large and reliably inducible (Fig. 1c). Since GIRK1^F137S^ appeared slightly faster in its activation time in comparison to the other subunits we decided to use GIRK1^F137S^ in all further constructs (Fig. 1d). However, depending on the experimental design and the specific cell-type or tissue, which shall be targeted such as brain, heart or gut, GIRK4^S143T^ might be more suitable to be used for co-expression in future experiments. The constructs have been designed in a way that allows modular access to exchange channel/opsin or tag in a singular cloning step to easily employ a different channel or sensor. GIRK-OpnX transfected HEK293 cells were readily modulated by light (Fig. 2), while solely G_i/o_-based tools were significantly faster in their activation times than the promiscuous G_i/o/q/11_ Opn4L. Overall GIRK-OpnX allows for a robust inhibition of cellular targets in HEK293 cells (Fig. 1-5), Purkinje cells (Fig. 6) and hiPSC-CMs (Fig. 7).

### Different light-activated GPCRs reveal differences in constitutive activity

GIRK-OpnX constructs displayed basal/constitutive inward currents under high potassium conditions (Fig. 3). This basal GIRK current may involve the constitutive activity of the respective GPCR or the GIRK channel itself. GIRK channels are directly activated by the binding of Gβγ subunits that dissociate from G protein heterotrimers upon GPCR activation. This causes a conformational change that opens the channel pore and allows a potassium flux^56–58,60,61,120^. Constitutive activity of G_i/o-_coupled GPCRs will therefore always activate a certain amount of GIRK channels. The lower this constitutive activity, the stronger the effect of GIRK channel activation by GPCRs, since the maximum potassium current is limited by the amount of activatable GIRK. Basal potassium currents by GIRK pore mutants have been shown in *Xenopus* oocytes for the co-expression of GIRK1^F137S^ or GIRK4^S143T^ with muscarinic M2-receptors^88,89^ and in HEK293 cells co-expressing GIRK1^F137S^ with mGluR2 and mGluR7^121^. We observed differences in resting potassium currents in HEK293 cells (Fig. 3) but saw no change in baseline currents in acute slice recordings between the different opsins. Opn4L activates both the G_q/11_ and the G_i/o_ pathway (Fig. 5 & 6). This in itself does not seem to interfere with basal GIRK channel activity, as the basal GIRK activity was comparable between Opn4L and Opn1LW (Fig. 3b). However, Opn4L-GIRK1^F137S^ expression in HEK293 GIRK1, 2 cells showed suppressed calcium responses in comparison to HEK293 GIRK1, 2 cells that expressed Opn4L alone (Fig. 5). At the same time GIRK1^F137S^ enhanced light-induced GIRK currents (Fig. 5). Interestingly baseline currents were also significantly larger when GIRK1^F137S^ was expressed in GIRK1, 2 expressing cells indicating the influence of subunits on basal GIRK currents.

The GIRK1 C-terminus has strong affinity for Gβγ^122^ and significantly increases basal GIRK conductance^123^. Differences in basal currents between GIRK-OpnX constructs therefore may not necessarily reflect dark activity of opsins but their capacity to compete for Gβγ with GIRK1.

Basal, as well as agonist-dependent activity of GIRK is increased by activation of protein kinase C-ε (PKCε), which involves the phosphorylation of GIRK subunits leading to an increased affinity of the channel to the membrane phospholipid phosphatidylinositol-4,5-bisphosphate (PIP_2_)^124^. However, PIP_2_ depletion and PKC activation by G_q/11-_coupled GPCRs inhibit GIRK channel activity in heterologous expression systems. Whether PKC activates or inhibits GIRK channel activity seems to be isoform specific^125^. It has been shown in cardiomyocytes that acetylcholine mediated potassium currents can be enhanced or inhibited dependent on the stimulation of PKC isoforms present^126^. Both protein kinase A^127^ and PLC signaling through PIP_2_ and PKC^124,126,128^ regulate cardiac GIRK channels, also in particular by phosphorylation of GIRK1 C-termini^129^. A constitutive increase in basal GIRK current in the heart (acetylcholine-regulated potassium current) has been observed in patients with chronic atrial fibrillation because of abnormal PKC function^130^. Moreover, changes in GIRK channel activity (either increases or decreases) have been associated with impairments in hippocampal-dependent cognitive functions^131^. Therefore, depending on the cell-type, physiological and behavioral paradigms increasing or decreasing the basal activity of GIRK channels might be preferable. For robust G_i/o_ mediated neuronal silencing, a low basal and large ligand- or light-induced GIRK current is preferable unless a steady inhibition is the desired outcome. Thus, engineering or finding new GPCRs with low or high constitutive activity might be advantageous for opto- and chemogenetic applications in particular with the aim of application in disease treatment.

### Co-expression of GIRK shifts the pathway predominance of Opn4L from G_q/11_ to G_i/o_

Melanopsin is expressed in intrinsically photosensitive retinal ganglion cells (ipRGCs), where it, among other cell types and tissues, is suggested to induce Gα_q/11, 14_ signaling^95,96,113,132–134^. In HEK293 cells Opn4 also couples to α_i/o_^95,96^. An α_s_^135^ activation has been suggested, while cross-motive signaling (G_q_ activation of AC)^134^ and signaling outside the Gα_q_-family^136^ has been proposed for ipRGCs. We have used Opn4L to control G_q/11_ mediated signals in the dorsal raphe nuclei^137^, the visual cortex^138^ and the cerebellum^96,138^. In fact, expression and light-dependent activation of Opn4L in cerebellar Purkinje cells of mice leads to a G_q/11_ mediated increase in action potential firing *in vitro*^96,138^ and *in vivo*^96,138^. This is also demonstrated in this study, when Opn4L is expressed alone (Fig. 6b_1_, c). Co-expression of GIRK1^F137S^ and Opn4L shifts the G_q/11_ mediated increase in AP firing towards a G_i/o_-GIRK mediated decrease in AP firing of cerebellar PCs (Fig. 6b_2_, c). Similarly, co-expression of GIRK was needed in hiPSC-CMs to allow G_i/o_ mediated suppression of the spontaneous beating rate (Fig. 7), while a G_q_ mediated increase in Opn4L expressing cardiomyocytes could not be detected. This can be explained by the transient transfection resulting in rather low expression rate and the chance that we did not stimulate the pacemaking cells. The calcium clock machinery effect depends on the Na^+^/Ca^2+^ exchanger transport leading to a low amount of depolarization^139^, especially compared to the massive outward directed current generated by GIRK.

Opn4L activates G_i/o_ and G_q/11_ pathways simultaneously in HEK293 cells, but the mechanisms underlying the co-activation and dominance of different signaling pathways *in vivo* remain elusive. The majority of GPCRs couple to multiple G protein subtypes with different coupling efficiencies and kinetics^140,141^. It has been suggested that G proteins have a selective barcode, which interacts with the GPCR through distinct residues^16^, while over 90% of G_q_ and G_s_ coupling receptors are suggested to be promiscuous^141^. Large Gα-binding pockets, leading to binding of different G protein α subtypes have been proposed as an explanation^141^. How one receptor can simultaneously activate different intracellular signaling pathways and how specificity is modulated has to be investigated.

### Application of GIRK-OpnX constructs in basic research and gene therapy

GPCRs that couple to G_i/o_ proteins and activate GIRK channels play a crucial role in regulating neuronal excitability and neurotransmission in the brain^142^ and heart^143^. GIRK channels contribute significantly to the potassium conductance in neurons and help to maintain the resting membrane potential^142^. Disruption of G_i/o_ mediated GIRK channel function can therefore result in neuronal hyperexcitability, which is implicated in conditions such as mood disorders^78,144^ and epilepsy^145,146^, while neurodegenerative diseases often involve changes in GPCR signaling^75–78,142,144^. Brain regions that are majorly affected by the process of aging (frontal cortex, hippocampus, substantia nigra and striatum) suffer particularly from a loss in protein density, which is suggested for almost half of the G_i/o_-coupled GPCRs^147^. This is linked to alterations in activity that result in an increased vulnerability towards neurodegenerative diseases in the aging brain^147^. For example, G_i/o_-coupled dopamine D2 receptors regulate GIRK channels in the striatum^148^. Dysfunction in this pathway contributes to motor symptoms of Parkinson’s disease^149^ and is also involved in schizophrenia^150,151^. Mutations in GIRK channels^152^ or G_i/o_-coupled receptors like GABA_B_ can lead to seizures and epilepsy^153–155^, which is in general affected by the aforementioned hyperexcitability. Furthermore, alteration in G_i/o_ signaling can contribute to addiction mechanisms including (but not limited to) opioid^156–160^ and cannabinoid^161,162^ receptor signaling. In the heart, GIRK channels are activated by acetylcholine released from the vagus nerve, leading to hyperpolarization of cardiac pacemaker cells. GIRK-mediated inhibition slows the heart rate, making GIRK channels crucial for regulating cardiac excitability and heart function^112^. Therefore, mutations in GIRK channels^163^ or G_i/o_-coupled receptors like the muscarinic M2-receptor can cause and alter arrhythmias^164^. Altered GIRK channel function due to increased G_i_ or decreased β-adrenergic receptors also impairs cardiac contractility and contributes to congestive heart failure progression^165^.

Thus, controlling G_i/o_ mediated GIRK channel activity, using co-expression cassettes with light-activated GPCRs or designer receptors, may be a therapeutic strategy to modulate or restore the excitation - inhibition balance and alleviate associated symptoms and dysfunctions^144,166^.

## Methods

### Animals and AAV injection

All experiments were conducted with approval of the animal care committee of Nordrhein-Westfalen (LANUV; Landesamt für Umweltschutz, Naturschutz und Verbraucherschutz Nordrhein-Westfalen, Germany; AZ.81-02.04.2019.A228). Experiments were performed in 2- to 6-month-old male and female C57BL/6J mice (The Jackson Laboratory, stock nr. 000664). Animals were housed in ventilated cages at a 12h/12h day and night cycle with access to food and water ad libitum.

For AAV injection mice were anaesthetized with isoflurane (initial dose 5% isoflurane 1.1 l min^-1^ air flow followed by a constant 1.3-2.0% isoflurane maintenance) and placed in a stereotaxic frame (Narishige, Japan) on a heating pad for body temperature maintenance. Mice received carprofen (2 mg/kg, s.c.) and buprenorphine (0.1 mg/kg, s.c.) for analgesia. Lidocaine (20 mg/kg) was administered subcutaneously after the surgery. Virus was pressure administered at two injection sites in a 1 µl dose at a steady flow rate in custom glass micropipettes (−7 mm AP, ±2 mm ML and 2-3 mm DL). Virus was expressed for 7 to 10 days for *in vitro* slice recordings while mice were individually housed and received postoperative care.

### Creation of plasmids

The cDNA for mouse codon optimized versions of GIRK1, 2 and 4 subunits (GIRK1 GenBank: AAH22495.1, GIRK2 NCBI Reference Sequence: NP_002231.1, GIRK4 NCBI Reference Sequence: NP_000881.3) was synthesized (GeneArt Gene Synthesis, Thermo Fisher Scientific) and subcloned into a custom vector variant of pAAV CW3SL eGFP (GenBank: KJ411916.2), which allowed the expression of GIRK subunit variants under control of a human cytomegalovirus (CMV) immediate early enhancer and CMV immediate early promoter but without a fluorescent label. The enhanced regulatory sequence CW3SL^119^ was included to improve transgene expression while reducing packaging size. Additionally, a vector was created that allowed the expression of GIRK2 alongside GIRK1 in the same reading frame but separated by the 2A peptide from porcine teschovirus-1 polyprotein (p2A). The cDNA for the Mus musculus melanopsin isoform 1 (Opn4L; NCBI reference sequence: NP_038915.1), the human long-wave-sensitive opsin 1 (Opn1LW; NCBI reference sequence: NP_064445.1), a mouse optimized version of the Lethenteron camtschaticum parapinopsin (ppOpn, UniProtKB/Swiss-Prot: Q764P5) and the Danio rerio Opsin7b (Opn7b, NCBI Reference Sequence: NP_001303878.1) was subcloned into a custom vector based on pAAV CW3SL eGFP (GenBank: KJ411916.2). The respective cDNA was expressed under control of a CMV immediate early enhancer and CMV immediate early promoter and in fusion with enhanced green fluorescent protein^167^, while a codon optimized version of the F137S GIRK1 mutant^88^ (GIRK1; GenBank: AAH22495.1) was placed upstream in the same reading frame but separated by a p2A site (GIRK-OpnX).

### AAV production

Recombinant adeno-associated virus (AAV) was created using the AAV helper free system (Agilent Technologies). HEK293T cells, stably expressing E1, were transfected with a pAAV vector carrying the respective ITR flanked gene of interest, pAAV-RC2/1 delivering the AAV-2 replication and the AAV-1 capsid gene and pHelper for adenoviral genes E2A, E4 and VA using polyethylenimine (1:5) in DMEM. Cells and medium were harvested 72 h after transfection. After separation by centrifugation (300g, for 10 min, at 4 °C), the cell fraction was resuspended in lysis buffer (0.15 M NaCl in 50 mM Tris-HCl buffer) and further disrupted by freeze-thaw cycling before DNase treatment (0.13 mg/ml DNase I, 13 mM MgCl_2_ at 37°C for 30 min). The supernatant was incubated with polyethylene glycol (12.5% PEG 8000, 0.5 M NaCl, for 2h, at 4 °C) for precipitation. After centrifugation (3700g, for 20 min, at 4 °C) the precipitate was resuspended with the purified cell lysate and stored overnight at 4 °C. After PEG precipitation and centrifugation (12.5% PEG 8000, 0.5 M NaCl, 2h; 3700g, for 20 min, at 4 °C) for a buffer transfer to 50 mM HEPES 1:1 (v:v), chloroform was added for the purification of viral particles. After thorough vortexing, low speed centrifugation (370g, for 5 min, at room temperature) allowed for disruption and removal of non-viral contaminants, which collect in the organic layer. The aqueous phase was then filtered (0.22 µm) and once again precipitated with PEG (12.5% PEG 8000, 0.5 M NaCl, for 2h; 3700g, for 20 min, at 4 °C) before resuspending the pellet in 0.001%-Pluoronic F-68 in PBS. After brief centrifugation at 13000g AAVs were stored at −80 °C until use.

### Human embryonic kidney cell 293 culture

HEK293 cells were maintained in Dulbecco’s Modified Eagle’s Medium (4.5 g/l D-glucose, Sigma-Aldrich), supplemented with 10% FBS (Gibco), 1% penicillin-streptomycin (Gibco), and 1% G418 for HEK293T cells or 1.5 µg/ml puromycin for HEK293 Opn4L cells in a humidified incubator at 37 °C and 5% CO_2_ in the dark. Cells were routinely subcultered upon reaching ∼ 70-85% confluence and prepared for experiments at this time point. Cells were transfected by admixing FuGENE® HD (Promega) transfection reagent in a ratio of 3.5 µl : 1 µg DNA in 100 µl Opti-MEM (Gibco) 12 – 24h before recordings.

### Preparing hiPSC-CMs

Human cardiomyocytes were differentiated from induced pluripotent stem cell line UMGi014-A clone 2, which was created from peripheral mononuclear blood cells from a healthy male donor^168^. hiPSC-CMs differentiation and purification involved modulation of Wnt signaling and lactate-based metabolic selection as described previously^169^. Differentiated hiPSC-CMs were cryopreserved for further use. For measurements, 1.5 – 2 million cells were thawed and plated on a 6-well-plate coated with 16 mg/mL Geltrex (Thermo Fisher Scientific) and maintained in hiPSC-CMs culture medium (RPMI 1640 medium (GlutaMAX; Thermo Fisher Scientific), 2% B27 (Thermo Fisher Scientific) supplemented with 2 µM thiazovivin (Sigma-Aldrich) at 37 °C and 5% CO_2_. Medium was replaced after two days to remove residues of thiazovivin and changed every 2-3 days afterwards.

hiPSC-CMs were dissociated and plated on 10 mm diameter round coverslips in cardiac cell clusters comprising 50000 cells on top of a spatially limited area, coated with an 8 µl drop of Geltrex/DMEM/Ham’s F-12. Cells were incubated in this drop for 40 min to build cardiac cell clusters. Afterwards the medium was filled up with hiPSC-CMs culture medium. After two days cardiac cell clusters started to beat spontaneously. Medium was replaced every 2-3 days.

Transient transfection of hiPSC-CMs with pAAV CMV mOpn4L.eGFP CW3SL or pAAV CMV hGIRK1^F137S^ p2A mOpn4L.eGFP CW3SL was performed using Promega ViaFect Transfection reagent (Promega) in a ratio 4:1 (Transfection reagent:DNA) within the cardiac cell clusters, following manufacturers recommendations.

### Contraction and frequency analysis of cardiac cell clusters

Experiments with beating cardiac cell clusters were performed within 48 h after transfection. hiPSC-CMs were incubated for 1-2 h in a 24-multiwell plate with 5 µM 9-*cis* retinal (Sigma-Aldrich) in culture medium. Beating was recorded at room temperature using a UI-306xCP-M camera (Imaging Development Systems GmbH, Germany). Online analysis of contractions was done with the custom-made Myocyte online Contraction Analysis (MoCA) software^114^ (available at github: https://github.com/awagdi0/MOCA/) giving the average of motion vectors per frame |V| as output through NI 9263 CompactDAQ (National Instruments) recorded with Powerlab 8/35 and LabChart 8.1.16 software for analysis. Spontaneous beating was tested with 5 s long light pulses of 460 nm as supramaximal light effect at 2.7 mW/mm^2^.

The light effect was analyzed by comparing the number of beats within the 30 s before the light pulse to the time period of 30 s starting with the illumination. 460 nm light was generated by a LED (serial number 1711-57 (PSU:1707-165), filtered through a F39-479 (474/27) excitation filter and reflected onto the cells via a 561 nm beamsplitter (AHF analysetechnik AG, F38-561) within an inverted IX73 microscope and through a 20x objective (LUCPLFLN20XPH/0,45). Light intensity was calibrated with the S170C sensor and Powermeter (Thorlabs). Imaging light was filtered by a 780 nm long pass filter (Schott RG780) to avoid excitation of Opn4L and GIRK-Opn4L by the illumination.

### Histology for cardiomyocytes

Cardiac cell clusters were fixed in 4% formaldehyde for 20 min, permeabilised with 0.2% Triton X100 for 20 min and stained in DPBS (Sigma-Aldrich) supplemented with 5% donkey serum for 2 h with primary antibodies against cardiac troponin T (Thermofisher, MA1-24611; 1:800) and for 1 h with secondary antibodies conjugated with Cy5 (711 −175-151, JacksonLab; 1:400) diluted in DPBS with 1:1000 DAPI (0018860.01, Th.Geyer) at room temperature. Cardiac cell clusters were imaged with LSM 800 confocal microscope (Zeiss) equipped with spectral multi-alkali photomultiplier detectors. Z-stack acquisition was performed using a Plan-Apochromat 20x/0.8 M27 via the ZEN 2.6 software (blue edition, Zeiss) software at 1024 x 1024 pixel resolution, with a stack step of 1.5 µm. Each imaged line was averaged 4 times during acquisition. Settings for pinhole were of 30 µm for Dapi, 24 µm for GFP, and 40 µm to image Cy5. Pictures were post-processed using ImageJ Fiji^170^.

### Electrophysiology – HEK293

Growth medium was exchanged for an external solution (20 mM NaCl, 120 mM KCl, 2 mM CaCl_2_, 1 mM MgCl_2_, 10 mM HEPES, pH 7.3 (KOH)) immediately before recording. Whole-cell patch clamp recordings were performed utilizing custom glass pipettes (2 – 4 MΩ, MPC325, Sutter Instrument) filled with an internal patch solution (100 mM L-Aspartic acid potassium, 40 mM KCl, 40 mM MgATP, 10 mM HEPES, 5 mM NaCl, 2 mM EGTA, 2 mM MgCl_2_, 0.01 mM GTP, pH 7.3 (KOH)). Low potassium conditions were achieved by exchanging the external solution (138 mM NaCl, 2 mM KCl, 2 mM CaCl_2_, 1 mM MgCl_2_, 10 mM HEPES-NaOH, pH 7.3 (KOH)) with a pneumatic pump (PicoPump PV820 World Precision Systems). Currents under low potassium conditions were always recorded after recordings in high potassium solution. Both patch solutions were supplemented with 1 µM of 9 cis-retinal. Signals were filtered by a 10 kHz Bessel filter in series with a 2.9 kHz 4-pole Bessel filter with an USB amplifier (EPC10, HEKA) and monitored by PatchMaster (v2X52, HEKA). PatchMaster also controlled voltage and a monochromator (Polychrome V, Till Photonics). Induced currents were calculated as the difference between averaged baseline and peak currents, while the peak current was averaged in a ±100 ms time window around the peak. Voltage ramps were recorded by clamping the cell to −60 mV for 100 ms before a step to −120 mV, which was held for 50 ms before gradually increasing the voltage to 20 mV with a slew rate of 0.4 mV/ms. Inducible current responses were either assessed at −60 mV after a 10 s light stimulation, or as the difference between currents elicited during voltage ramps. Cells were stimulated with light during an initial ramp (total stimulus duration 600 ms) and then monitored for 20 additional ramps; the currents displayed are calculated as the difference between the initial and the 10th ramp. Kinetics were estimated by an exponential tail end fit (I_(t)_ = I_0_ + e^-t/τ^) during the light phase or in the dark for the calculation of τ_off_ for OPN1LW and Opn7b. Intensities were calculated by recording light with an optical power meter (PM100D with S142C, Thorlabs) set to the respective wavelength. The mean irradiance measured over the course of a minute was *E_360 nm_* = 0.058 mW/mm², *E_470 nm_* = 1.23 mW/mm², and *E*_560 nm_ = 0.87 mW/mm².

### Electrophysiology – acute cerebellar slice

All experiments were performed in red light. Acute sagittal, cerebellar slices (250 µm) were prepared 10-14 days after AAV injection (Vibratome VT1000S, Leica) in a dissection solution (87 mM NaCl, 2.5 mM KCl, 0.5 mM CaCl_2_, 7 mM MgCl_2_, 1.25 mM NaH_2_PO_4_, 25 mM NaHCO_3_, 10 mM D-Glucose and 75 mM sucrose enriched with 95% O_2_ and 5% CO_2_). Slices were incubated in recording solution (125 mM NaCl, 2.5 mM KCl, 2 mM CaCl_2_, 1 mM MgSO_4_, 1.25 mM NaH_2_PO_4_, 26 mM NaHCO_3_, and 20 mM D-Glucose enriched with 95% O_2_ and 5% CO_2_) at 37 °C for 45-60 min and then stored at room temperature until use. Slices were preincubated for 20 min in recording solution containing 25 µM 9-*cis*-retinal, 0.025% (±)-α-tocopherol (Sigma-Aldrich), and 0.2% essentially fatty acid free albumin from bovine serum (BSA, Sigma-Aldrich) before use. Custom glass pipettes (3 – 5 MΩ, MPC325, Sutter Instrument) were filled with internal solution: 125 mM Potassium Gluconate, 10 mM HEPES, 4 mM NaCl, 2 mM MgCl_2_, 0.2 mM EGTA, 4 mM Mg-ATP, 0.4 mM Na-GTP, and 10 mM Tris-Phosphocreatine (pH 7.3). For voltage-clamp recoding 5 µM lidocaine N-ethyl bromide (QX-314; Tocris Bioscience) were added to block Na^+^ currents and action potential generation. For recordings using the constitutive active Opn7b 10 µM ZD7288 (Sigma-Aldrich) were added to the external solution to block HCN-currents. Purkinje cells were visually identified using a 40x objective attached to an upright microscope (BX51WI, Olympus) under infrared illumination and a Polychrome V (TILL Photonics) set to 470 nm light was used to identify expressing cells. Whole-cell recordings were low pass-filtered at 3 kHz using an EPC10 amplifier (HEKA) and conducted on cells that were voltage clamped to −60 mV. Light dependent GIRK currents were measured by applying an increasing voltage ramp (−130 to −50 mV in 1 s, 3 repetitions) before and during light applications (ppOpn: 360 nm; Opn4L/ Opn7b: 470 nm; Opn1LW: 595 nm). Light application started 5 s before the ramp. To measure light induced and opsin mediated effects on neuronal activity the cells were held at a constant base current that elicited moderate spontaneous activity before a train of 10 s different light pulse conditions was applied (light off/470 nm/light off/560 nm/light off). To measure membrane potential, rheobase, spike rate and input resistance the response to different currents steps was recorded (−200 to 300 pA, step 50 pA, 1 s current step, 2 s repetition rate) PatchMaster (v2X52, HEKA) was used for data acquisition and MATLAB (The MathWorks, Inc.) for offline analysis.

### Fluorescence imaging

Calcium signals reflecting G_q/11_ pathway activation were elicited in HEK293 cells constitutively expressing GIRK1, 2 subunits and transfected with CMV Opn4L.eGFP or CMV GIRK1^F137S^ p2A Opn4L.eGFP constructs alongside CMV GCaMP6f CW3SL by blue light stimulation 470 nm (*E_470 nm_* = 1.23 mW/mm², Polychrome V, Till Photonics). All monitored cells were recorded while in whole cell configuration (see above, external solution: 20 mM NaCl, 120 mM KCl, 2 mM CaCl_2_, 1 mM MgCl_2_, 10 mM HEPES, pH 7.3 (KOH), internal solution: 125 mM Potassium Gluconate, 10 mM HEPES, 4 mM NaCl, 2 mM MgCl_2_, 0.2 mM EGTA, 4 mM Mg-ATP, 0.4 mM Na-GTP, and 10 mM Tris-Phosphocreatine, pH 7.3). Currents and fluorescence were recorded at the same time. Images were taken at an inverted microscope (Axiovert 35, Carl Zeiss) with an exposure time of 100 ms and a sCMOS camera (Prime BSI Express, Photometrics) controlled by Micromanager 2^171^/ Image J (ImageJ 1.53c, NIH). Time series analyzer v3 plugin was used for analysis of calcium signals over time. The fluorescence signal was background corrected and is displayed as relative change in fluorescence normalized to the average of the first three data points (0.3 s) at the beginning of the recording (ΔF/F0). The maximum fluorescence intensity displayed, reflects the intensity of individual cells at the timepoint where the average of cells reaches its maximum intensity, but is averaged within ±0.5 to control for fluctuations in the signal.

### Histology

Viral expression was assessed in sagittal sections of C57BL/6J mice after an expression period of 14 days. After lethal dosage of ketamine (100 mg/kg, i.p.) and xylazine (20 mg/kg, i.p.) mice were transcardially perfused with phosphate buffered saline (PBS, pH 7.4) followed by 4% paraformaldehyde (PFA, Sigma Aldrich). Post fixation ensued for 2-8 h in PFA. Brains were cryopreserved by immersion in 30% sucrose (Fischer Chemicals) until loss of buoyancy. After freezing at −20 °C in tissue tek (Sakura) specimen were sagitally sectioned (35-40µm) using a Cryostat (Leica CM3050 S). Slices and cells were assessed at a confocal microscope (Leica TCS SP5II, LAS AF 2.6, Leica Microsystems) using a 63x/1.4 oil objective (Excitation: 488 nm, HCX PL APO CS) with an excitation wavelength of 480 nm to visualize eGFP.

### Statistics and post-processing

Statistical analysis and the generation of graphs was conducted using SigmaPlot 12.5 (Systat Software GmbH), JASP 0.16.4.0 (Jasp Team) or custom written Matlab (The MathWorks Inc.) scripts. Figures were created using CorelDRAW2018 (Corel Corporation) and ImageJ Fiji^156^ (Fiji 2.1.0/ ImageJ 1.53c, NIH). Displayed Images of cells have been inverted and post processed for visual clarity. Frequency changes between cardiac cell clusters were analyzed by one-way ANOVA with Tukey’s multiple comparison procedure. Other statistical differences are displayed as the results of analysis of variance (ANOVA) followed by Holm-Sidak multiple comparison procedure or (paired) t-test if the data were normally distributed (Shapiro-Wilk p > .05) and showed equal variance (p > .05), else differences were tested for by Kruskal-Wallis ANOVA followed by Dunn’s multiple comparison procedure or Mann-Whitney rank sum or Wilcoxon signed rank sum test. Individual *p*-values are displayed in the respective figure while the full test statistics can be found in supplementary table 1.

## Acknowledgements

This work was supported by funds from Gamut Therapeutics (now SparingVision) and by Deutsche Forschungsgemeinschaft (DFG) grants: Project ID 316803389 - SFB 1280 (S.H.); HE 2471/21-1, (S.H.): Project number 492434978 - GRK 2862/1, Subproject (01 (S.H.; 09 (I.S.)). Further support came from the German Research Foundation through the Cluster of Excellence (EXC2067) Multiscale Bioimaging and by the German Centre for Cardiovascular Research (DZHK) (L.C. and T.B.).

## Author Contributions

Conceptualization: S.H., I.S., T.B., D.D. and B.M.. Experimentation: I.S., A.G., L.C. and B.M.. Data analysis: I.S., A.G. and B.M.. Writing – original draft: S.H., A.G. and B.M.. Writing - review & editing: S.H., I.S., D.D. A.G., L.C., T.B. and B.M.. Funding acquisition: S.H., T.B. and I.S..

---

**SUPPLEMENTARY TABLE 1:** STATISTICAL PROCEDURES AND COMPARISONS.

| **Figure** | **Test** | **Comparison** | **Test-statistic** | **P-value** |
| --- | --- | --- | --- | --- |
| **1c** | **Kruskal-Wallis** | Current density, functional channels | $H(3, 54) = 35.73$ | **<.001** |
| | Dunn's | GIRK1F137S vs. GIRK4S143T | $Z = 0.4296$ | **>.999** |
| | Dunn's | GIRK1F137S vs. GIRK2 | $Z = 5.586$ | **<.001** |
| | Dunn's | GIRK1F137S vs. GIRK1, 2 | $Z = 2.184$ | .174 |
| | Dunn's | GIRK4S143T vs. GIRK2 | $Z = 4.707$ | **<.001** |
| | Dunn's | GIRK4S143T vs. GIRK1, 2 | $Z = 1.610$ | .644 |
| | Dunn's | GIRK2 vs. GIRK1, 2 | $Z = 3.067$ | **.013** |
| **1d** | **Kruskal-Wallis** | $\tau$ on, all | $H(3, 55) = 14.248$ | **.003** |
|  | Dunn's | GIRK1F137S vs. GIRK4S143T | 3.69 | **.001** |
|  | Dunn's | GIRK1F137S vs. GIRK2 | 1.881 | .360 |
|  | Dunn's | GIRK1F137S vs. GIRK1, 2 | 2.210 | .163 |
|  | Dunn's | GIRK4S143T vs. GIRK2 | 1.728 | .504 |
|  | Dunn's | GIRK4S143T vs. GIRK1, 2 | 1.359 | >.999 |
|  | Dunn's | GIRK2 vs. GIRK1, 2 | 0.3447 | >.999 |
| **1e** | **Kruskal-Wallis** | $\tau$ off, all | $H(3, 55) = 1.403$ | .252 |
| **2c** | **Kruskal-Wallis** | $\tau$ on, all | $H(2, 53) = 16.004$ | **<.001** |
| | Dunn's | Opn4L vs Opn1LW | $Z = 3.049$ | **0.007** |
| | Dunn's | Opn4L vs ppOpn | $Z = 3.967$ | **<.001** |
| | Dunn's | Opn1LW vs ppOpn | $Z = 1.1$ | .814 |
| **2d** | **ANOVA** | $\tau$ off, all | $F(2, 53) = 2.747$ | .073 |
| **2f1** | **Paired t** | Opn4L, dark vs 470 nm | $t(12) = 9.331$ | **<.001** |
| **2f2** | **Wilcoxon Signed Rank** | Opn1LW, dark vs 560 nm | $W = -210$ | **<.001** |
| **2f3** | **Paired t** | ppOpn, dark vs 360 nm | $t(22) = 10.609$ | **<.001** |
| **2f4** | **Paired t** | Opn7b, dark vs 560 nm | $t(9) = -7.012$ | **<.001** |
| **3a1** | **Wilcoxon Signed Rank** | Opn4L, $K^+$ vs 470 nm | $W = -61$ | **.033** |
| **3a2** | **Paired t** | Opn1LW, $K^+$ vs 560 nm | $t(20) = 6.395$ | **<.001** |
| **3a3** | **Wilcoxon Signed Rank** | ppOpn, $K^+$ vs 360 nm | $W = -231$ | **<.001** |
| **3a4** | **Paired t** | Opn7b, $K^+$ vs 560 nm | $t(12) = -6.495$ | **<.001** |
| **3b1&2** | **ANOVA** | Contribution (%) to current | $F(2, 52) = 6.958$ | **.002** |
| | Holm-Sidak | Opn4L vs Opn1LW | $t = 0.741$ | .462 |
| | Holm-Sidak | Opn4L vs ppOpn | $t = 2.964$ | **.005** |
| | Holm-Sidak | Opn1LW vs ppOpn | $t = 3.333$ | **.009** |
| **4e** | **Paired t** | Opn4L, dark vs 470 nm | $t(5) = -1.687$ | .152 |
| **4e** | **Paired t** | GIRK-Opn4L, dark vs 470 nm | $t(5) = 2.704$ | **.043** |
| **4f** | **Paired t** | Opn1LW, dark vs 595 nm | $t(6) = 0.063$ | .952 |
| **4f** | **Wilcoxon Signed Rank** | GIRK-Opn1LW, dark vs 595 nm | $W = -21$ | **.031** |
| **4g** | **Paired t** | ppOpn, dark vs. 360 nm | $t(7) = 1.541$ | .167 |
| **4g** | **Paired t** | GIRK- ppOpn, dark vs 360 nm | $t(6) = 4.348$ | **.005** |
| **4h** | **Paired t** | Opn7b, dark vs 470 nm | $t(7) = 0.451$ | .451 |
| **4h** | **Wilcoxon Signed Rank** | GIRK-Opn7b, dark vs 470 nm | $W = 45$ | **.004** |
| **5** | **Mann-Whitney Rank Sum** | Light-induced calcium, Opn4L vs GIRK-Opn4L | $U = 19$ | **0.043** |
| **5** | **t** | Light-induced GIRK current, Opn4L vs GIRK-Opn4L | $t(17) = -2.530$ | **0.022** |
| **6c** | **t** | $\Delta$ spontaneous AP, Opn4L vs GIRK-Opn4L | $t(15) = 4.093$ | **<.001** |
| **6e** | **Wilcoxon Signed Rank** | $\Delta$ evoked AP, Opn4L dark vs 470 nm | W = 91 | **<.001** |
| **6f** | **Paired t** | $\Delta$ evoked AP, GIRK-Opn4L dark vs 470 nm | t(9) = 2.596 | **.029** |
| **6g** | **Mann-Whitney Rank Sum** | $\Delta$ MP, Opn4L vs GIRK-Opn4L | U = 82 | .900 |
| **6h** | **Mann-Whitney Rank Sum** | $\Delta$ rheobase, Opn4L vs GIRK-Opn4L | U = 13 | **.008** |
| **6j** | **Paired t** | Input resistance, Opn4L dark vs 470 nm | t(15) = 0.894 | .386 |
| **6j** | **Wilcoxon Signed Rank** | Input resistance, GIRK-Opn4L dark vs 470 nm | W = -34 | **.016** |
| **7c** | **ANOVA** | Frequency change | F (2, 22) = 30.11 | **<.001** |
|  | Tukey<br>Tukey<br>Tukey | control vs. Opn4L<br>control vs. GIRK-Opn4L<br>Opn4L vs. GIRK-Opn4L | p-value corrected for multiple comparisons: | **<0.001**<br>**<0.001**<br>0.5015 |

